# Supra-threshold perception and neural representation of tones presented in noise in conditions of masking release

**DOI:** 10.1101/575720

**Authors:** Katharina Egger, Torsten Dau, Bastian Epp

## Abstract

The neural representation and perceptual salience of tonal signals presented in different noise maskers were investigated. The properties of the maskers and signals were varied such that they produced different amounts of either monaural masking release, binaural masking release, or a combination of both. The signals were then presented at different levels above their corresponding masked thresholds and auditory evoked potentials (AEPs) were measured. It was found that, independent of the masking condition, the amplitude of the P2 component of the AEP was similar for the same stimulus levels above masked threshold, suggesting that both monaural and binaural effects of masking release were represented at the level of P2 generation. The perceptual salience of the signal was evaluated at equal levels above masked threshold using a rating task. In contrast to the electrophysiological findings, the subjective ratings of the perceptual signal salience were less consistent with the signal level above masked threshold and varied strongly across listeners and masking conditions. Overall, the results from the present study suggest that the P2 amplitude of the AEP represents an objective indicator of the audibility of a target signal in the presence of complex acoustic maskers.

## Introduction

One major task of the auditory system is to distinguish between different sound sources and to segregate single sources from the acoustic background. For sound segregation and identification, the auditory system utilizes monaural cues, such as spectro-temporal information contained in the sound, as well as binaural cues, such as interaural disparities occurring at the two ears (i.e.,interaural level- and time differences). The ability of the auditory system to benefit from these cues is often assessed via the effect of masking of a tonal signal by a noise masker. Many aspects of auditory signal detection in noise can be accounted for by the power spectrum model of masking [1].This model assumes that the frequency selectivity of the auditory system can be represented as a bank of overlapping band-pass filters and that the detection threshold of a tonal signal in the presence of a noise masker is mainly determined by the signal-to-masker energy ratio at the output of the band-pass filter centered at the signal frequency.

Some masking data, however, cannot be accounted for by the power spectrum model. For example, when the masker exhibits coherent intensity fluctuations across frequency, the masked thresholds of signals presented in noise are lower than predicted by the power spectrum model. This monaural phenomenon of improved signal detectability due to coherent intensity fluctuations of the masker across frequency has been referred to as comodulation masking release (CMR) (for a review, see [2] and [3]). There are two experimental CMR paradigms, the band-widening paradigm and the flanking-band paradigm. In the band-widening paradigm, the masker is spectrally centered at the signal frequency and is either unmodulated or comodulated. The bandwidth-dependent difference in masked threshold between the comodulated and the unmodulated masker conditions is then referred to as CMR, or modulated-unmodulated difference [4]. In the flanking-band paradigm, the masker consists of several narrow bands of noise whereby one masker band is centered at the signal frequency, the signal centered band (SCB), and one or more bands, the flanking bands (FBs), are spectrally separated from the signal frequency (e.g., [5–7]). In this paradigm, CMR has been defined in two ways: 1) as the difference in masked threshold between the conditions with the SCB alone and when the SCB and comodulated FBs are presented; or 2) as the difference in masked threshold for unmodulated versus comodulated masker bands.

A release from masking can also be found when the signal and the masker are presented at different spatial locations compared to a condition in which both sounds are presented at the same location. Differences in the spatial location of the signal and the masker lead to differences in binaural cues between the signal and the masker. These binaural cues can lead to an enhanced signal detectability which is commonly referred to as binaural masking level difference (BMLD) [8–9]. Effects of BMLD have typically been investigated in experiments with tonal signals masked by noise where either the signal or the noise masker contained an interaural phase difference (IPD)[10]. The BMLD has been defined as the difference in threshold between the diotic condition, where the signal and the masker are presented in phase across ears, and a dichotic condition, where either the signal or the masker is presented interaurally out of phase.

The effects of CMR, BMLD, and their combination have commonly been studied at signal levels close to masked threshold (e.g., [11–16]), whereas their contribution to the perception of signals above masked threshold (i.e., supra-threshold) has received much less attention. The benefit of CMR at supra-threshold levels has mainly been discussed in connection to either speech detection and recognition tasks (e.g., [17–19]) or regarding signal intensity and frequency discrimination tasks(e.g., [20–21]). In terms of binaural processing, the perceived intensity of a signal, i.e., its loudness, has primarily been studied in conditions of BMLD at different supra-threshold levels of the signal in the presence of noise (e.g., [10,22–23]). More recently, Verhey and Heise [24] investigated the magnitude of the tonal content as well as the partial loudness of a tonal signal in the presence of a noise masker. In their study, conditions with and without monaural and binaural masking releases at different signal levels above the respective masked thresholds were used. Verhey and Heise [24] showed that the masking releases were strongest at low signal levels above masked threshold and decreased towards higher levels. The above studies were all either focused on monaural processing, regarding CMR, or on binaural processing, regarding BMLD, whereas the supra-threshold perception of the combined effect of CMR and BMLD has not yet been investigated. It is therefore still unknown to what extent the two effects contribute to supra-threshold perception.

Various physiological studies investigated the neural mechanisms underlying the processing and representation of the auditory cues that lead to a perceptual masking release. A physiological correlate of CMR in human listeners has been proposed at cortical levels using electroencephalography (EEG) [25], magnetoencephalography (MEG) [26], as well as functional magnetic resonance imaging (fMRI [27–28]). Physiological correlates of BMLD in humans have also been investigated at different levels along the auditory pathway using EEG [29–31], MEG [32–33], and fMRI [34].

Billings et al. [35] compared late auditory potentials evoked by a speech syllable presented in speech-shaped noise with the performance in a behavioral sentence identification task at different signal-to-masker ratios. They identified the N1 component of the electrophysiological response as a correlate of behavioral performance. However, while the N1 component was found to be mainly dependent on the signal-to-masker ratio, the behavioral data showed a signal-level dependent behavior, complicating the relation between behavioral performance and evoked potentials amplitude.

Epp et al. [36] recorded evoked potentials of masked tonal signals in conditions of masking release at different signal-to-masker ratios. At a fixed signal-to-masker ratio, the level of the signal above masked threshold was altered by the introduction of monaural across-frequency cues to the masker (yielding CMR), binaural cues to the tonal signal (yielding BMLD), or a combination of both. Epp et al. [36] found a combined effect of CMR and BMLD reflected in the N1/P2 complex of the evoked potential. The N1 component was sensitive to the introduction of binaural cues, but did not show any sensitivity to monaural across-frequency cues. In contrast, the amplitude of the P2 component at a given signal-to-masker ratio was found to be larger in conditions with comodulated noise than in conditions with uncorrelated noise. A larger P2 amplitude was also observed when binaural cues were introduced to the tonal signal as well as when both cues were present. In fact, the P2 amplitude was of similar size in experimental conditions where the signal was presented at a similar level above its masked threshold. This was not the case for the N1 component, nor the P2-N1 difference amplitude. Epp et al. [36] therefore suggested that the P2 component might be a potential candidate for the neural representation of the partially masked signal at the corresponding level above masked threshold.

The hypothesis of the present study was that signals at the same level above masked threshold evoke the same P2 amplitudes in the corresponding evoked potential. It was further postulated that there exists a perceptual correlate of the signal level above masked threshold, the “perceptual salience”, and that signals at the same level above masked threshold evoke the same perceptual salience. To test this, the present study extended the investigations of Epp et al. [36] and considered three experiments: In the first experiment, behavioral detection thresholds of masked tonal signals were obtained in conditions reflecting either purely monaural, purely binaural, or a combination of monaural and binaural masking releases. In a second experiment, AEPs for the masked signals were recorded in the same listeners at individually adjusted signal levels corresponding to equal levels above their respective masked thresholds. Finally, in a third experiment, the listeners rated the perceptual salience of the signal embedded in the noise masker at equal signal levels above their respective masked thresholds.

## Materials and methods

### Stimuli

The signal was a 700 Hz tone of 300 ms duration, including 20 ms raised-cosine on- and offset ramps. The signal was presented either with the same interaural phase, i.e., an IPD of 0 degree (diotic condition), or interaurally out of phase (dichotic condition) with an IPD of 150 degree^1^.

The masker consisted of either five narrowband noise bands (“multi-band conditions”) or one broadband noise (“broadband condition”). In the multi-band conditions, each noise band was 24 Hz wide and had a sound pressure level (SPL) of 50 dB. One noise band (SCB) was centered at the signal frequency (700 Hz) and the other four bands, the FBs, were arranged symmetrically (on a linear frequency scale) with respect to the SCB at 300, 400, 1000, and 1100 Hz. The remote spectral distance between the SCB and the FBs was chosen in order to reduce within-channel cues contributing to CMR (e.g., Cohen, 1991). The noise bands had either uncorrelated or comodulated intensity fluctuations across frequency, subsequently denoted as UN and CM masker, respectively. The broadband (BB) masker was an 824-Hz wide noise band centered at the signal frequency (700 Hz) and had an overall level of 60 dB SPL. This level was chosen since it produces approximately the same amount of energy that passes the auditory filter centered at the signal frequency as the SCB in the multi-band conditions [37]. Hence, a similar masking effect produced by the BB masker was expected as for the UN masker.

All maskers were presented diotically and had a duration of 900 ms, including 20 ms raised-cosine on- and offset ramps. The signal was presented in the last 300 ms of the masker. This was done to avoid a temporal overlap of the masker onset response and the signal onset response in the AEPs. The signal in both binaural conditions was combined with each of the maskers resulting in six different stimulus conditions: the diotic signal presented in either uncorrelated noise (UN_0_), comodulated noise (CM_0_), or broadband noise (BB_0_) and the dichotic signal presented in the same masker types (UN_150_, CM_150_, and BB_150_, respectively).

The noise bands were generated by multiplying a random-phase sinusoidal carrier at the desired frequency with a low-pass noise without a DC component. The low-pass noise was generated in the frequency domain by assigning numbers between ±0.5 from a uniformly distributed random process to the real and imaginary parts of the respective frequency components. For the UN and BB maskers, independent realizations of the low-pass noise were used for each masker band, while the same low-pass noise was used for all five bands of the CM masker. The masking noise was newly generated for each interval and each trial.

### Apparatus

All stimuli were digitally generated in MATLAB with a sampling rate of 44100 Hz and a 16-bit resolution, converted from digital to analogue (RME DIGI96/8 PAD, experiments 1 and 3; RME Fireface UCX, experiment 2) and presented via circumaural headphones (Sennheiser HD580, experiments 1 and 3) or via in-ear headphones (Etymotic ER-2, experiment 2). In experiment 2, the Psychophysics Toolbox extensions for MATLAB [38–39] were used. The headphones were calibrated and equalized at the signal frequency of the tone.

AEPs were recorded using a BioSemi ActiveTwo measurement system. The listeners wore an elastic cap with plastic electrode holders for 64 sintered Ag/AgCl pin-type electrodes. The electrode holders were filled with highly conductive, Signa electrode gel to reduce the contact impedance between electrode and skin. The common reference was the electrode placed at the left mastoid (P9), according to the extended 10/20 layout as standardized by the American Electroencephalographic Society (e.g., [40–41]). The electrodes were connected to the ActiveTwo AD-box, which amplified and performed A/D conversion of the measured potentials with a sampling rate of 1024 Hz and a 24-bit resolution. The potentials were recorded with the data acquisition software ActiView (version 6.05), which streamed the continuous EEG to hard disk. During the recordings, an anti-aliasing digital low-pass filter with a cut-off frequency of 200 Hz was used together with a digital high-pass filter with a cut-off frequency of 0.16 Hz to reduce the influence of slow, non-neural potentials, such as skin potentials [42].

### Listeners

Eight listeners (three female, five male), aged between 22 and 28 years, participated in the experiments. None of them reported any history of hearing impairment. All listeners had pure-tone hearing thresholds within 15 dB HL for the standard audiometric frequencies from 125 to 4000 Hz. All listeners were paid an hourly wage for their participation. The listeners were the same in all three experiments. During the psychoacoustical experiments, the listeners were seated in a double-walled, sound-attenuating booth. The AEP experiment was carried out in a double-walled and electrically shielded, sound-attenuating booth. All participants provided informed consent and all experiments were approved by the Science-Ethics Committee for the Capital Region of Denmark (reference H-KA-04149-g).

### Procedure

In the first experiment, detection thresholds of the masked signals were measured for each listener in all six stimulus conditions (UN_0_, CM_0_, BB_0_, UN_150_, CM_150_, and BB_150_). An adaptive, three interval, three-alternative forced-choice procedure with visual feedback was used. The intervals within a trial were separated by pauses of 500 ms. The listeners had to indicate the interval in which the signal was presented by pressing the corresponding key on the keyboard. The adaptive signal level adjustment followed a one-up two-down algorithm to estimate the 70.7% point of the psychometric function [43]. The initial step size was 8 dB. After each lower reversal, the step size was halved until it reached the minimum step size of 1 dB. This step size was kept constant for another six reversals of which the arithmetic mean was calculated and used as the estimated threshold for that run. Each listener performed four threshold measurements per stimulus condition whereof the arithmetic mean of the last three runs was taken as the final individual threshold estimate. The different stimulus conditions were presented in random order within blocks, where each condition occurred once.

In the second experiment, late AEPs for the masked signals were recorded for all stimulus conditions. The signal levels were adjusted to 10, 15, 20, and 25 dB above the masked thresholds of the individual listener (obtained in experiment 1). The stimuli were presented in random order, evenly distributed over 400 sweeps (i.e., 400 repetitions per stimulus condition and level). The presentations were separated by an inter-stimulus interval of 300 ms plus an additional jitter of 250 to 350 ms. During the EEG recordings, the listeners were presented a quiet movie of their choice with subtitles on a low-radiation screen. The listeners were asked to relax but not to fall asleep and to avoid moving as far as possible. The experiment was divided in six blocks of approximately 35 minutes each, whereof two blocks were recorded in one session. The listeners participated in three sessions distributed over different days.

In the third experiment, the listeners rated the perceptual salience of the tone embedded in the masking noise for all stimulus conditions by magnitude estimation. The signal levels were individually-adjusted to −10, 0, 5, 10, 15, and 20 dB relative to the masked thresholds. The test signals to be judged (i.e., the tones embedded in masking noise) were presented in paired trials with a fixed reference which was a diotic 700-Hz tone presented at 70 dB SPL. In each trial, the reference tone and the test signal could be played as often as the listeners desired and were accessed either by pressing the corresponding keys or by clicking the corresponding buttons with the mouse. The number of times the reference and the test signal were presented in each trial was recorded. The listeners had to rate the salience of the tone in the test signal on an endpoint-anchored scale by using a continuous slider. The scale was anchored by ‘not audible’ on the lower end and by ‘reference’ on the upper end. These anchors were combined with a scale without labels in between, as suggested in [44]. The slider’s position was mapped onto a numerical scale between 0 (designating the lower anchor ‘not audible’) and 10 (designating the upper anchor ‘reference’). The listeners were not aware of the mapping. Prior to the experiment, the listeners were told that the experiment contained the same types of stimuli they had been listening to in the previously performed measurements.

To familiarize the listeners with the task, a short training session was provided before the initial run of the experiment, using stimulus examples of varying perceptual saliences. Four different types of stimulus examples were presented: (1) the reference tone; (2) the masking noise alone, randomly representing either the UN, CM, or BB masker; (3) the masked signal in one stimulus condition randomly selected out of all six (UN_0_, CM_0_, BB_0_, UN_150_, CM_150_, or BB_150_), at a supra-threshold level, corresponding to 25 dB above masked threshold; and (4) randomly selected stimuli of all stimulus conditions with signal levels as used in the experiment.

The stimuli were tested in random order within blocks, where each level for each stimulus condition occurred once. Sixteen blocks were presented, whereof the very first block was considered as additional training and excluded from further data analysis. The individual salience ratings were therefore derived from fifteen responses per stimulus condition and level.

### Data analysis

The continuous EEG data were processed off-line in MATLAB to extract the late AEPs. Single sweeps of a time interval of 1200 ms (150 ms pre-stimulus period and 1050 ms following the stimulus onset) were extracted corresponding to the different stimulus conditions and levels. For each sweep, a linear fit was subtracted from the waveform to remove low-frequency potential drifts (e.g., [45]). The fit was based on the intervals 150 ms before the signal onset (at 600 ms) and 150 ms after the stimulus. The data were then digitally filtered with a zero-phase, second order forward-backward Butterworth low-pass filter with a cut-off frequency of 20 Hz. Each sweep was baseline corrected by subtracting the arithmetic mean over the 150 ms pre-stimulus period. An iterative weighted averaging method was used to average all single sweeps per stimulus condition and level for each listener [45]. The weighted average was computed by weighting each single sweep by the inverse power of its noise [46]. Single sweeps which exceeded the artefact rejection threshold of ±100 μV in any of the recorded channels were discarded and not included in the average. This resulted in the average AEP including the response to the masker onset.

To obtain the neural response following the signal onset within the masker for each listener, referred to as the AEP change complex, the AEP in the latency interval between 600 and 1050 ms was extracted from the averaged sweeps. Baseline correction was applied considering a 150 ms pre-stimulus period (with respect to the signal onset at 600 ms). The grand mean AEP change complex was computed as an arithmetic mean over all individual AEPs.

In addition, the amplitude components of the AEP change complex were evaluated individually for each listener. The change complex can be described by the response extrema N1, measured between 90 and 190 ms, and P2, measured between 180 and 290 ms (with respect to the signal onset). Peak amplitudes for the N1 and P2 components were extracted for the electrode placed at the vertex (Cz), individually for each listener. Electrode Cz was used since it showed the largest response averaged across all conditions. Peak amplitudes were extracted separately for each stimulus condition and level, using a semi-automatic procedure including identification of N1 and P2 components by a peak-scan-algorithm and subsequent verification by visual inspection of all waveforms. The algorithm located minima and maxima in the time windows defined for N1 and P2, respectively. The peaks were indicated automatically if the first derivative of the potential showed a zero-crossing and had a change in sign within the time window. If multiple extrema were found, the algorithm selected the ones with the largest amplitude. If an extremum with the wrong polarity or no response maximum was found by the algorithm, peaks were selected manually by requiring a clear peak within the time window with correct curvature.

The behavioral salience data were analyzed in MATLAB using a non-parametric Friedman test. The effects of level and stimulus condition on the rated signal salience were tested. The analysis was based on median salience ratings across all listeners. Since no interaction effects can be tested using the Friedman test, two separate tests were conducted. To test the effects of level, the data obtained at the different levels were arranged in columns and the stimulus conditions were represented along the rows of the data matrix. To test the effects of stimulus condition, the matrix was transposed before testing.

## Results & Discussion

### Experiment 1: Masked signal detection thresholds in noise

Masked thresholds for the eight individual listeners as a function of the signal IPD are shown in Fig 1 (A-H). The error bars in each panel indicate the standard deviations of the thresholds. The bottom right panel shows the grand average thresholds across the listeners. The results are shown for the UN masker (squares), the CM masker (triangles), and the BB masker (circles). For all masker types, the thresholds obtained in the dichotic conditions (150 degree IPD) were lower than those obtained in the diotic conditions (0 degree IPD), reflecting a BMLD. All listeners, except for the one represented in panel A, showed a decrease in threshold when the masker envelope across frequency was changed from being uncorrelated (UN masker) to comodulated (CM masker), reflecting a CMR effect. This was the case in both binaural conditions. For listener A, the difference in masked threshold between the UN and the CM maskers, i.e., the effect of CMR, nearly vanished in the dichotic condition. The thresholds obtained with the signal presented in the BB masker were similar to those obtained with the UN masker, except for listener A, who showed a considerably lower threshold for the BB masker in the diotic condition than all other listeners. The amount of BMLD obtained for the different masker types differed across the listeners. The BMLD ranged from 8.7 dB (listener H) to 20.4 dB (listener A) for the UN masker, from 7.4 dB (listener H) to 14.6 dB (listener F) for the CM masker, and from 8 dB (listener A) to 13 dB (listeners B and D) for the BB masker. The amount of CMR decreased with increasing IPD for half of the listeners (A, C, D, and E). Three listeners (B, G, and H) showed roughly the same amount of CMR for the two signal IPDs; listener F showed a larger CMR in the dichotic condition compared to the diotic condition. The diotic CMR varied from 9 dB (listener E) to 12 dB (listener C) and the dichotic CMR varied from 1.2 dB (listener A) to 12.4 dB (listener F).

**Fig 1.**
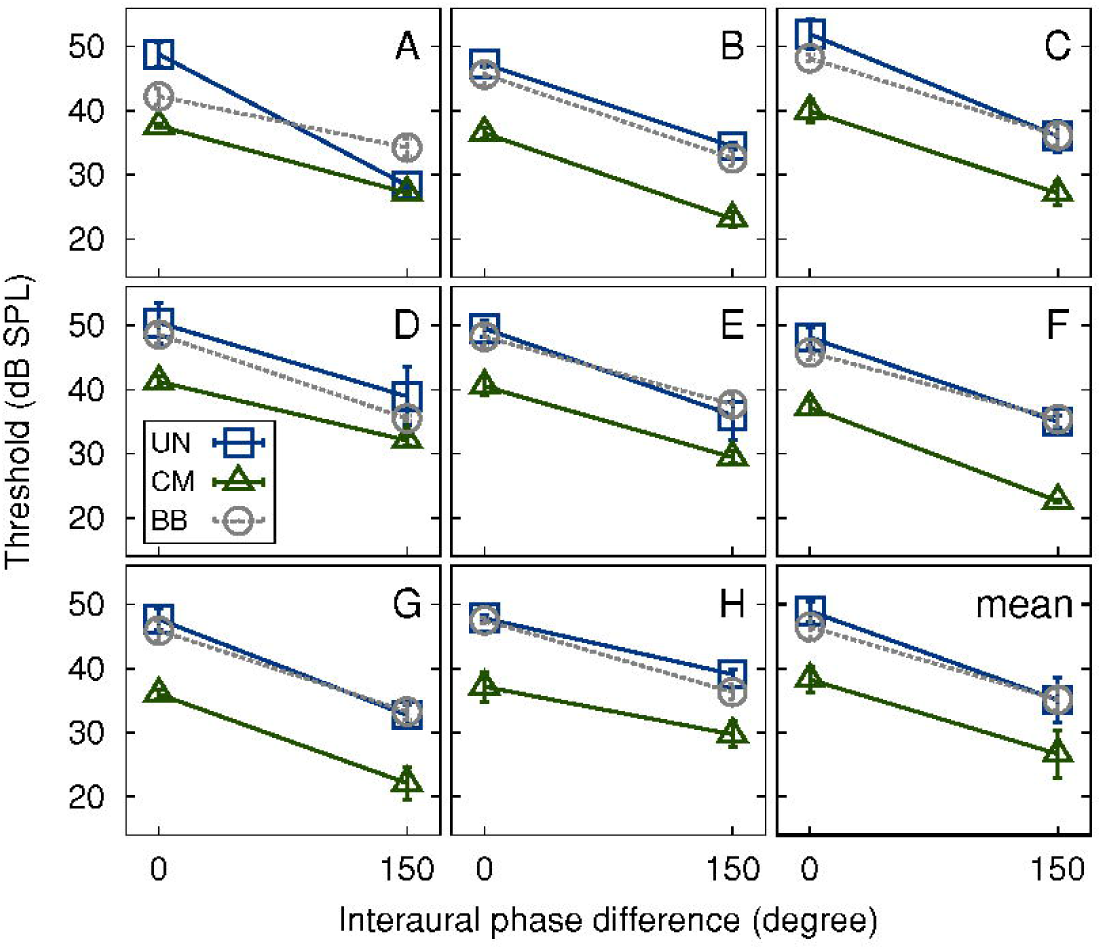
Mean thresholds. Individual mean thresholds of the listeners (panels A-H) and grand average thresholds across all listeners (bottom right panel) as a function of signal IPD. Thresholds are plotted for UN (squares), CM (triangles), and BB (circles) maskers. Error bars indicate plus-minus one standard deviation.

Regarding the average data, a benefit was observed by introducing masker comodulation (yielding CMR), a signal IPD (yielding BMLD), or a combination of both. The thresholds were largest for UN_0_ (48.9 dB) and smallest for CM_150_ (26.7 dB). CMR reached a magnitude of 10.7 dB in the diotic condition and decreased to 8.4 dB in the dichotic condition. Thresholds were lower in the dichotic conditions than in the diotic conditions, demonstrating a BMLD of 13.9 dB in the case of the UN masker and approximately 11.5 dB in the cases of the CM and BB maskers.

The data are consistent with findings reported in the literature [13,16,36]. Interestingly, also in Epp and Verhey [13] and Hall et al. [16], a few listeners showed a strong reduction of CMR in the dichotic relative to the diotic condition in combination with large BMLDs, whereas the majority of the listeners showed a smaller BMLD and a comparable CMR in the diotic and dichotic conditions.

### Experiment 2: Auditory-evoked potentials to tones in conditions of masking release

The grand mean AEPs across all listeners, for the latency interval of 550 to 950 ms relative to the stimulus onset, i.e., the interval associated with the AEP change complex, is shown in Fig 2. The three panels illustrate the results obtained with the diotic signals (solid lines) and the dichotic signals (dashed lines) in the presence of the UN masker (left panel), the CM masker (middle panel), and the BB masker (right panel), respectively. The ordinate represents different levels of the signal above the individual listeners’ masked thresholds. Independent of the masker condition (UN, CM, and BB) and the binaural condition (0 versus 150 degree IPD), the magnitude of the AEP change complex increased with increasing signal level relative to masked threshold (bottom to top in Fig 2). This is consistent with the initial hypothesis of this study that a higher signal level above masked threshold yields a stronger neural response. However, for a given level above masked threshold, the grand mean AEP magnitude was larger in the dichotic (dashed) than in the diotic (solid) conditions. Hence, the grand mean data, when ignoring possible individual differences in latency of the AEP, is at odds with the initial hypothesis of this study that signals presented at equal levels above masked threshold evoke the same AEP magnitude, independent of the stimulus condition.

**Fig 2.**
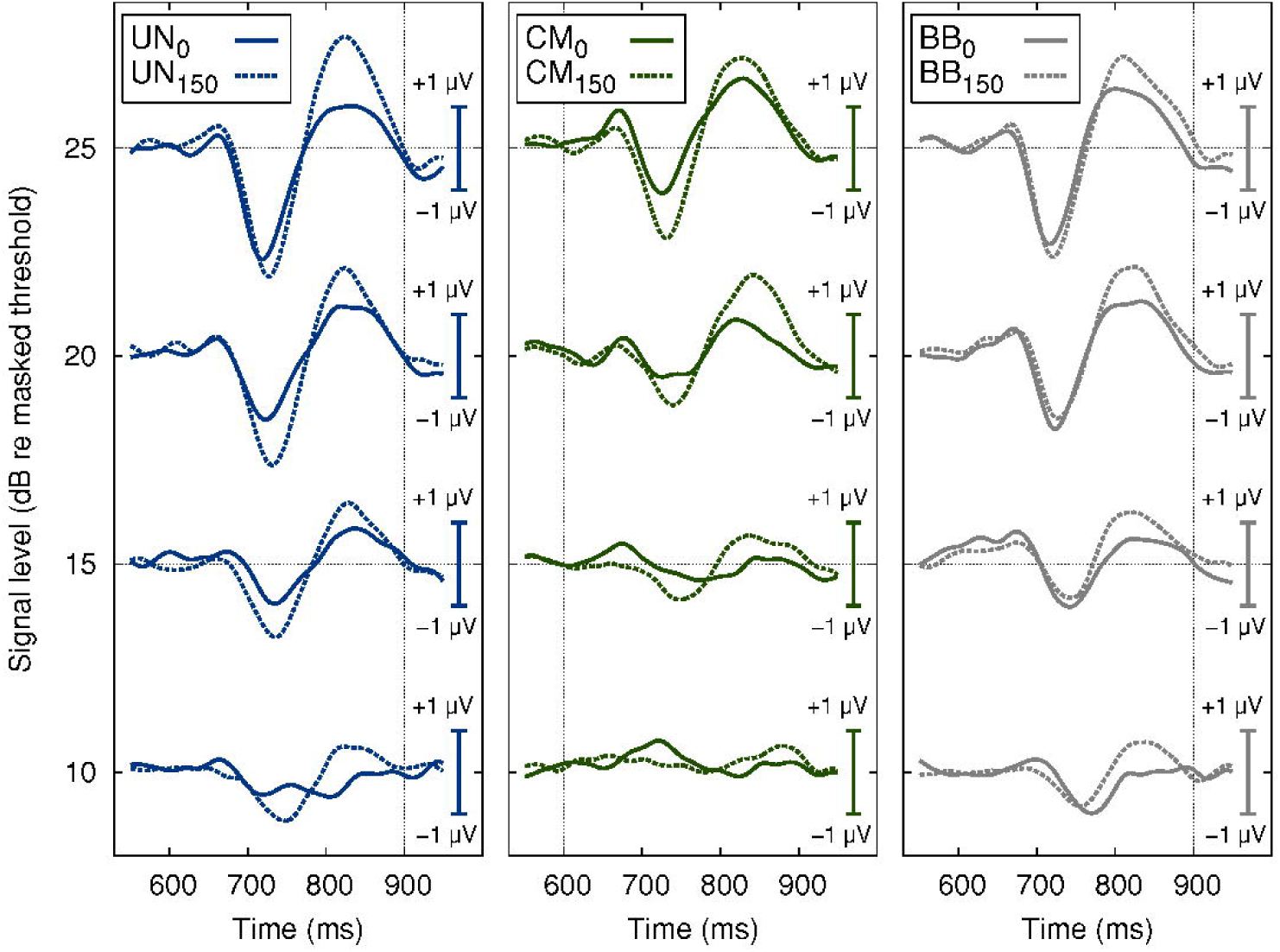
Grand mean AEP. Grand mean evoked potential data for all listeners for a latency interval of 550 to 950 ms relative to the stimulus onset. The three panels illustrate the averaged AEPs measured for diotic (solid lines) and dichotic (dashed lines) signals in the presence of UN (left), CM (middle), and BB (right) maskers. Waveforms are plotted for four different signal levels: 10, 15, 20, and 25 dB relative to masked thresholds (indicated on the y-axes on the left-hand side). The potentials were baseline corrected considering a 150 ms pre-stimulus period (with respect to the signal onset at 600 ms).

The results shown in Fig 2 are compatible with the data obtained in the study of [47] who measured AEPs evoked by a diotic or antiphasic tone in a broadband masker. In their study, larger responses were found for antiphasic than for diotic signals at a fixed level above masked threshold. However, only the N1-P2 peak of the change complex was evaluated and only three listeners were considered. The grand average AEP data show also a similar trend as the results obtained in Epp et al. [36], where larger grand average potentials were obtained for dichotic than for diotic signals. In the study by Epp et al. [36], the signal was however presented at equal SPLs rather than at equal levels above masked threshold.

The grand average AEPs, however, do not account for individual differences in latency across listeners which might challenge the interpretation of the grand mean AEP. To compensate for individual differences in AEP latency, the peak amplitudes of the change complex were extracted and analyzed individually for each listener^2^. The grand mean of the extracted N1 (left panel), P2 (middle panel), and P2-N1 (right panel) amplitudes as a function of the signal level relative to the individual listeners’ masked thresholds are shown in Fig 3. The open symbols represent the stimulus conditions with the diotic signal in the presence of the UN (squares), CM (triangles), and BB (circles) maskers, respectively. The filled symbols represent the corresponding stimulus conditions with the dichotic signal. The magnitude of the N1 (negativity) increased with signal level. For a given level, the mean N1 amplitude varied between 1.1 and 2.2 µV across the stimulus conditions. The magnitude of the P2 (positivity) also increased with signal level but showed a markedly smaller amount of variability (of 0.5 to 1.1 µV) across stimulus conditions than N1. The difference amplitude, P2-N1, often considered in AEP studies, showed variability of up to 3 µV.

**Fig 3.**
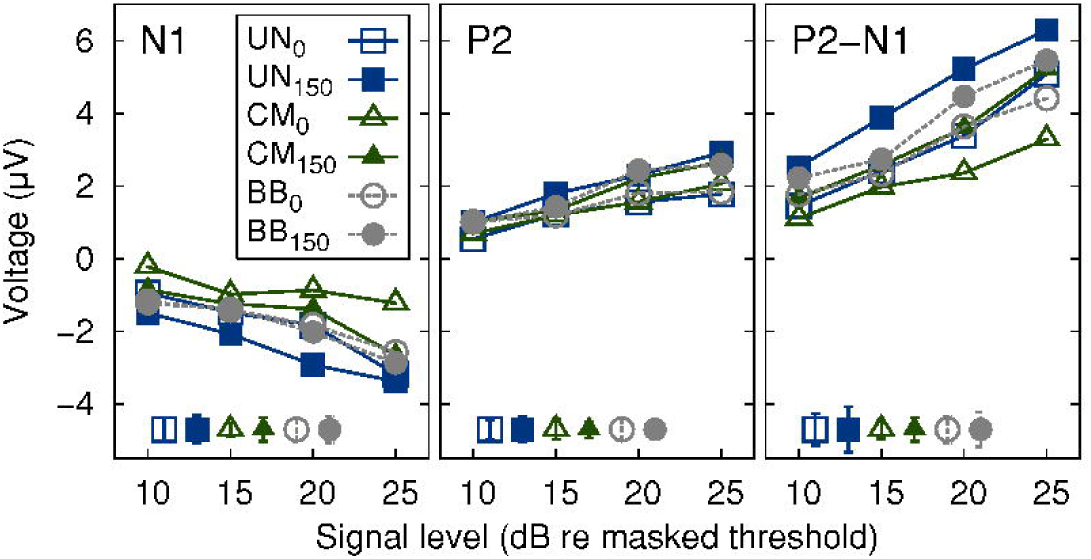
Mean amplitudes of AEP. Grand mean of amplitudes N1 (left panel), P2 (middle panel), and the difference between P2 and N1 (right panel) across all listeners as a function of signal level relative to masked threshold. Amplitudes are plotted for diotic (open symbols) and dichotic (filled symbols) signals in the presence of UN (squares), CM (triangles), or BB (circles) maskers. Error bars in the lower, left corner indicate plus-minus one standard error (averaged over all signal levels) for each stimulus condition, respectively.

The P2 amplitudes for both binaural conditions were found to be similar at equal levels above masked thresholds (Fig 3, middle panel), in agreement with the initial hypothesis of this study. Hence, individual differences in latency resulted in a broadening of the grand mean AEP peaks and therefore in different amplitudes for different conditions in Fig 2. The N1 amplitude (and, hence, the P2-N1 difference) showed a large variability across stimulus conditions and, in contrast to the P2 amplitude, no clear pattern between N1 amplitude and condition was found.

In summary, the present results support the hypothesis that the P2 component of the AEP change complex correlates with the signal level above masked threshold. Specifically, P2 may represent an objective indicator of the overall perceptual masking release, obtained here with monaural and binaural signal-masker configurations.

### Experiment 3: Perceptual salience ratings of tones in conditions of masking release

Panels A-H of Fig 4 show the individual salience ratings of the signal as a function of the signal level relative to the masked threshold. Since the salience scale had fixed endpoints, the data are represented by the median and the corresponding interquartile ranges. The open symbols represent the results for the diotic conditions, whereas the filled symbols indicate the results for the dichotic conditions obtained with the UN masker (squares), the CM masker (triangles), and the BB maskers (circles), respectively. For all listeners, the salience ratings increased with increasing signal level in each stimulus condition. The salience tended to be highest for stimulus conditions without a release from masking (UN_0_ and BB_0_). Two listeners (F, G) rated the salience of the dichotic tone embedded in the CM masker (CM_150_) clearly lower than in all other stimulus conditions. Some listeners (A, C, D, and H) showed a similar increase in salience with increasing signal level relative to masked threshold, particularly for the CM masker conditions. Other listeners (B, E, F, and G) showed ratings that are ordered according to their physical signal-to-masker level ratio rather than according to the signal level relative to masked threshold. The salience for signals that were presented below masked threshold (i.e., −10 dB SL) nearly vanished for all but listeners A and C.

**Fig 4.**
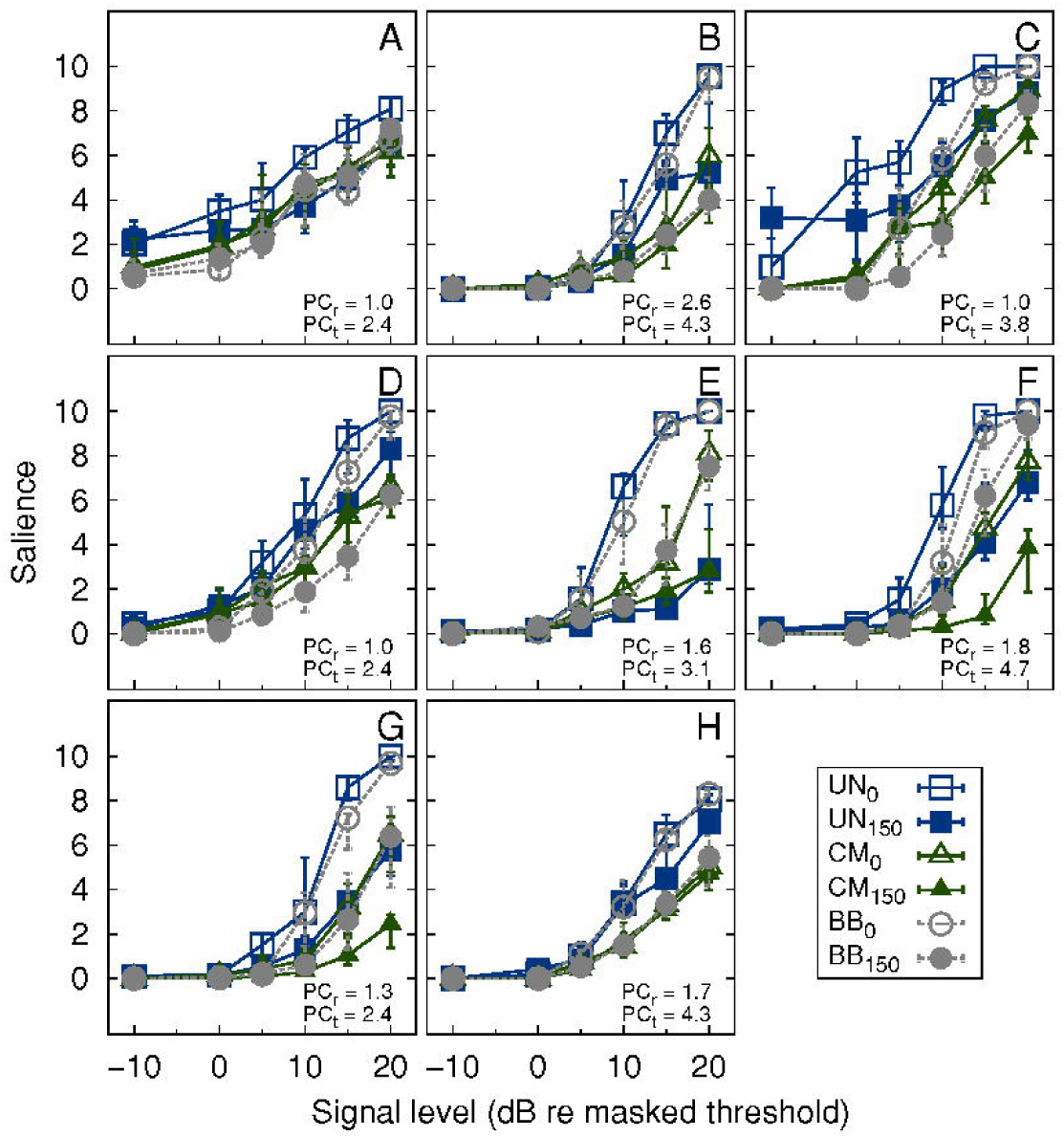
Individual salience ratings. Individual median salience ratings of the individual listeners (panels A-H) as a function of masked threshold. Salience is plotted for diotic (open symbols) and dichotic (filled symbols) signals in the presence of UN (squares), CM (triangles), and BB (circles) maskers. Error bars indicate the interquartile range (panels A-H). PC_r_ and PC_t_ refer to the average number of times the reference and the test signal were presented in each trial, respectively.

A non-parametric Friedman test was used on the pooled data to investigate the effects of level and stimulus condition on the rated salience. The test provided a significant main effect of level [chi^2^(5) = 224.03, p < 0.001] and of stimulus condition [chi^2^(5) = 71.03, p < 0.001]. The significant effect of stimulus condition is contrary to the initial hypothesis of this study that signals at the same level above masked threshold evoke the same perceptual salience for all stimulus conditions.

The variability across listeners observed in these data suggests that different perceptual attributes might have influenced the salience rating. The perception of a tone in the presence of a background noise has been discussed mainly in connection to perceptual attributes like tonality, tonalness, pitch strength, and partial loudness, or in relation to measures of annoyance (for a review, see [48]). Effects of monaural and/or binaural masking releases on tone perception have not been considered in this context yet. The order of the ratings at a given signal level (above threshold) in one of the listener groups (B, E, F, G) suggests that the partial loudness of the signal might have contributed to the judgments.

### Relation between auditory evoked potential amplitude and salience rating

The salience ratings increased monotonically with increasing signal level relative to masked threshold. However, for a given level above masked threshold, the rated salience differed considerably across stimulus conditions and showed a large variability across listeners. These results are not consistent with the AEP data for the P2 component and, thus, do not support the initial hypothesis of this study that masked signals evoking similar P2 amplitudes result in the same perceptual salience of the signal. The statistical significance of stimulus condition further provides evidence against the validity of this hypothesis.

The observed mismatch between the salience ratings and the electrophysiological responses using the measurement paradigm chosen in the present study may be caused by the dominance of certain perceptual attributes in the salience rating. The individual contributing factors, such as partial loudness or tonalness, would need to be investigated further by isolating them separately in additional perceptual experiments.

An attempt to relate partial loudness to electrophysiological data was made by Rupp et al. [49]. They measured auditory evoked fields (AEFs) in response to stimulation with tone pulses in the presence of Schroeder-phase tone-complex maskers and compared the AEFs with patterns of a loudness experiment obtained with the same listeners. While the tone pulses were kept at a constant sound pressure level, the temporal position of the pulses within the maskers was varied, resulting in varying neuromagnetic response amplitudes and a varying partial loudness of the tone pulses. The behavioral measure in Rupp et al. [49] was a two-alternative-forced choice paired-comparison of the stimuli. The paired comparisons were then projected on a perceptual ratio scale using the Bradley-Terry-Luce (BTL) method [50]. Rupp et al. [49] found a correlation between the AEFs and the partial loudness patterns. In particular, in those conditions where the tone pulses were masked more effectively by the tone-complex maskers, the corresponding AEF showed a smaller amplitude than in the conditions where the signal was masked less effectively. The BTL method has also been used to measure other perceptual attributes of sounds like annoyance with subsequent analysis of other contributing factors [51]. Since loudness most likely was a contributing factor to the perceptual salience of the signal in the present study, a similar approach might help to clarify the observed disparity between AEP amplitude and salience rating.

## Summary and conclusions

The present study investigated the neural representation and perceptual salience of tonal signals presented in different noise maskers. The maskers and the target signals were chosen such that they produced different amounts of either monaural masking release (CMR), binaural masking release (BMLD), or a combination of both. The signals were then presented at different levels above their respective masked thresholds. AEPs were used as an objective indicator of the signal’s neural representation in the maskers. It was found that signals presented at equal levels above the listeners’ individual masked threshold (but at different physical intensities) resulted in similar amplitudes of the P2 component (but different amplitudes of the N1 component) across various conditions of masking release. This suggests that both monaural and binaural effects of perceptual masking release are represented at the level of P2 generation. In contrast to the electrophysiological findings, the subjective ratings of the signal salience showed a significant effect of stimulus condition. Hence, the perceptual salience as considered in the present study did not show a consistent correlation with the corresponding P2 amplitude patterns. The reason for this might be the contribution of several other perceptual cues to the salience ratings, with relative weights that may vary across listeners. Possibly, the partial loudness and the magnitude of tonal content of the supra-threshold tone embedded in the complex masker contributed to the perceived salience of the signal.

Overall, if the correlation of the P2 amplitude of the AEP change complex with the level above masked threshold also holds true for other conditions of masking release and more complex signals like speech, P2 might serve as an indicator of the supra-threshold signal audibility in the presence of a complex masker in various conditions of perceptual masking.

## Acknowledgments

We thank all our listeners for their participation in this study.

## Footnotes

1 An IPD of 150 degrees was encouraged by the maximum IPD that human listeners can perceive due to the limitation of the circumference of the head, based on the average maximum, nonambiguous IPD calculated out of head-related transfer functions (HRTFs) of 68 listeners (HRTF database of the Acoustics Research Institute, Austrian Academy of Sciences)

2 Even though the latency was shown to represent an informative dimension in the evaluation of AEPs (Billings et al., 2009), this dimension is closely correlated to the amplitude of the change complex component and shows the same or an even larger variability.

